# Size-Based Characterization of Freshwater Dissolved Organic Matter finds Similarities within a Water Body Type across Different Canadian Ecozones

**DOI:** 10.1101/2020.04.03.024174

**Authors:** Pieter J. K. Aukes, Sherry L. Schiff, Jason J. Venkiteswaran, Richard J. Elgood, John Spoelstra

## Abstract

Dissolved Organic Matter (DOM) represents a mixture of organic molecules that vary due to different source materials and degree of processing. Characterizing how DOM composition evolves along the aquatic continuum can be difficult. Using a size-exclusion chromatography technique (LC-OCD), we assessed the variability in DOM composition from both surface and groundwaters across a number of Canadian ecozones (mean annual temperature spanning −10 to +6 C). A wide range in DOM concentration was found from 0.2 to 120 mg C/L. Proportions of different size-based groupings across ecozones were variable, yet similarities between specific water-body types, regardless of location, suggest commonality in the processes dictating DOM composition. A PCA identified 70% of the variation in LC-OCD derived DOM compositions could be explained by the water-body type. We find that DOM composition within a specific water-body type is similar regardless of the differences in climate or surrounding vegetation where the sample originated from.

**Highlights:**

- Size-exclusion chromatography (using LC-OCD) is a fast and effective tool to quantify differences in DOM composition across different environments
- Proportions of biopolymers and low molecular weight fractions can distinguish between surface and groundwater DOM
- Similar water-body types have comparable DOM size compositions across ecozones that range in annual air temperatures from –10 to 6ºC

## 1 INTRODUCTION

Dissolved organic matter (DOM) is a ubiquitous component of terrestrial and aquatic ecosystems. DOM influences light penetration within lakes (Schindler et al. 1996) and provides an energy source for microbial metabolism (Biddanda and Cotner 2002). Comprised of thousands of molecules with differing structures and properties, DOM concentration and composition can vary greatly among environments due to different physical, chemical, and biological processes. Future drinking water treatment options may be significantly impacted as climate change is predicted to alter the quantity and quality of DOM in surface waters (Ritson et al. 2014). Increased terrestrial DOM contributions observed among northern surface waters are thought to result in the ‘brownification’ of these systems, affecting lake characteristics and food webs (Creed et al. 2018; Wauthy et al. 2018). Hence, changes to DOM composition and concentration must be monitored or observed to better understand how a changing composition can impact subsequent water treatment effectiveness and downstream ecosystems.

DOM concentration and composition evolves along the aquatic continuum (i.e. transport pathway from headwaters to the ocean) on both global and local scales (such as along a river) due to additional DOM inputs, microbial and photosynthetic production, flocculation, and photochemical degradation (Massicotte et al. 2017; Ward et al. 2017; Xenopoulos et al. 2017). These differences in DOM are found between water body types (i.e. lake vs groundwater), as well as across an environmental gradient, due to the range in watershed characteristics, such as geology, hydrologic flow paths, vegetative communities, water residence time in the water body, and the duration of exposure to sunlight (Mueller et al. 2012; Jaffé et al. 2012; Kellerman et al. 2014; Xenopoulos et al. 2017). Determination of the variability in DOM composition across the aquatic continuum can indicate potential avenues of future DOM change, allowing us to better anticipate water treatment requirements and costs. However, due to the inherent complexity and heterogeneity of DOM, a number of different techniques have been used to quantify compositional differences.

Many of the techniques for analyzing DOM composition measure either bulk characteristics or only a subset of all DOM molecules by capturing select components. Generally, the need for enhanced information on DOM composition and molecular moieties results in increased cost and complexity to implement (McCallister et al. 2018). Fortunately, comprehensive chemical characterization of DOM is not always required to make useful predictions on how DOM will behave in the environment. For instance, DOM composition has been assessed via molecular ratio assays (Hunt et al. 2000), light absorption (Weishaar et al. 2003), fluorescence (Jaffé et al. 2008), resin fractionation (Kent et al. 2014), and mass spectrometry (Kellerman et al. 2014; Hutchins et al. 2017). Bulk optical indices, although common due to the relative ease of analysis, only respond to compounds that absorb or fluoresce in a specific range of wavelengths and can include non-organic components in the matrix (Weishaar et al. 2003) or miss non-absorbing DOM components (Her et al. 2002). The ideal DOM characterization method, or combination of methods, would provide sufficient detail on the key aspects of DOM composition that control its fate and function, yet be analytically simple enough to be cost effective and practical for environmental applications.

Size exclusion chromatography (SEC) has been used to characterize freshwater DOM in lakes (Kent et al. 2014), rivers or streams (He et al. 2016), and groundwaters (Szabo and Tuhkanen 2010). Liquid Chromatography – Organic Carbon Detection (LC-OCD) is a simple, fast (<120 minutes per sample), and quantitative SEC method that separates DOM based upon hydrodynamic radii (Huber et al. 2011), characterizing both absorbing and non-absorbing DOM components. Studies using LC-OCD have focussed on wastewater and water treatment applications (Ciputra et al. 2010), with some data from agricultural and forested catchments in Germany (Heinz et al. 2015), and rivers in South Korea (He et al. 2016) and Spain (Catalán et al. 2017). LC-OCD-based DOM groupings provide a relatively easy measure of DOM size-based composition across a large suite of environments.

Light absorbing and fluorescing components of DOM vary in the environment (Jaffé et al. 2008, 2012) but few studies examine how non-light absorbing DOM components differ across areas with different climate and vegetation. Here we determine the effectiveness of using size-based groupings of DOM to quantify differences in DOM composition and, to our knowledge, present one of few studies that pairs DOM characterization of both groundwater and surface water across a gradient of climate and vegetation regimes. The objectives of this study are: 1) to determine whether differences in DOM composition, from various water-body types and ecozones (an ecosystem unit that groups similar areas of biodiversity (Marshall et al. 1999)), can be identified using different LC-OCD fractions, and 2) to quantify the degree of similarity in DOM composition among water-body types collected from different ecozones.

## 2 METHODS

### 2.1 Site Descriptions

Samples were collected in the summer months (June to August) across various Canadian ecozones (Marshall et al. 1999) that span a mean annual temperature from −9.8 to 6.5ºC, precipitation from 313 to 940 mm, and overlie continuous to no permafrost (Figure 1; Suppl. Table 1). Surface water samples were collected from the Boreal Shield (IISD Experimental Lakes Area (ELA)), Mixedwood Plains (agriculturally-impacted Grand River (GR)), Taiga Shield (Yellowknife (YW) and Wekweètì (WK)), and Southern Arctic ecozones (Daring Lake (DL); Suppl. Info Table 1). Groundwater, defined here as water collected beneath the ground surface, was sampled from various depths from the Boreal Shield (Turkey Lakes Watershed (TLW)) Mixedwood Plains (Nottawasaga River Watershed (NRW)), Atlantic Maritime (Black Brook Watershed (BBW)), Taiga Shield (YK, WK), and Southern Arctic (DL). Both NRW and BBW are areas of extensive agriculture. Shallow groundwater samples were collected from the Northwest Territories (deepest extent of active-layer in July or August, ~0.1-0.5 m.b.s; meters below surface) and TLW (0-7 m.b.s), while samples from NRW and BBW were collected at depths ranging between 1 and 30 m.b.s.

**Figure 1:**
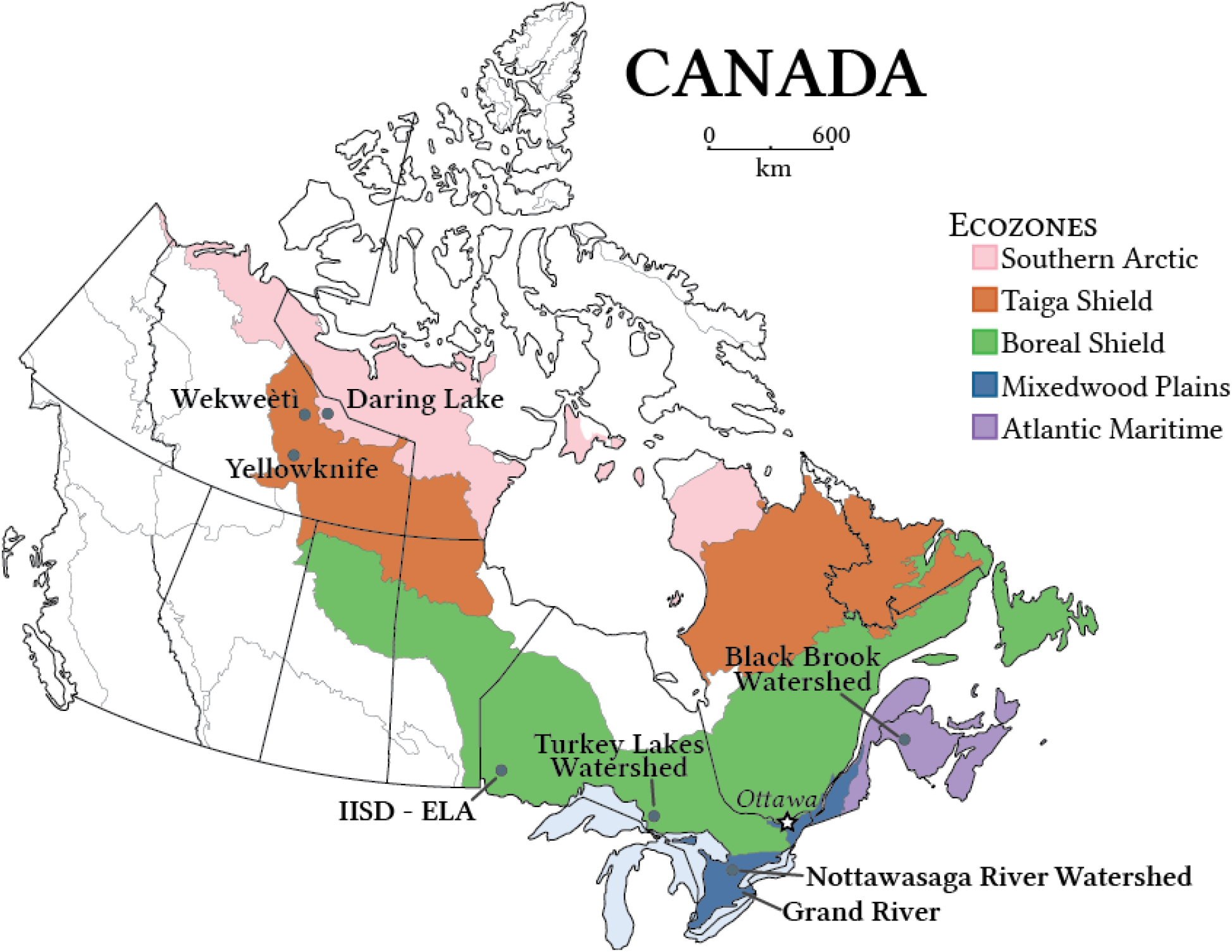
Ecozones (Marshall et al. 1999) and DOM collection sites: Daring Lake (DL; Southern Arctic ecozone), Wekweeti (WK; Taiga Shield ecozone), Yellowknife (YW; Taiga Shield ecozone), International Institute for Sustainable Development - Experimental Lakes Area (IISD-ELA; Boreal Shield ecozone), Turkey Lakes Watershed (TLW; Boreal Shield ecozone), Nottawasaga River Watershed (NRW; Mixedwood Plains ecozone), Grand River Watershed (GR; Mixedwood Plains ecozone), and Black Brook Watershed (BBW; Atlantic Maritime ecozone).

### 2.2 DOM Characterization

All samples were filtered to 0.45μm (Whatman GD/X) into acid-washed and pre-rinsed glass vials. Samples were immediately stored cool (<4°C) and dark until analyses (within two weeks). DOM composition was quantitatively assessed using LC-OCD (DOC-Labor, Karlsruhe, Germany; Huber et al. 2011) at the University of Waterloo. A description of analytical equipment and DOM fraction characterization has been described elsewhere (Huber et al. 2011) and a detailed explanation can be found in the Supplementary Information. Briefly, the sample is injected through a size-exclusion column (Toyopearl HW-50S, Tosoh Bioscience) that continuously elutes DOM based on hydrodynamic radii into five hydrophilic fractions (from largest to smallest): biopolymers (BP; polysaccharides or proteins), humic substances fraction (HSF; humic and fulvic acids), building blocks (BB; lower weight humic substances), low molecular weight neutrals (LMWN; aldehydes, small organic materials), and LMW-acids (LMWA; saturated mono-protic acids). Part of the sample bypasses the column for measurement of total dissolved organic carbon concentration, herein referred to as DOM concentration in mg C/L. As the hydrophilic component comprised 90±9% across all DOM samples, fraction percentages were normalized to the sum of the eluted components (total hydrophilic fraction, herein referred to as DOM_hyphl_). The LC-OCD is both calibrated and run with potassium hydrogen phthalate, IHSS-HA, and IHSS-FA standards to check DOM concentrations, minimize differences in machine drift, and compare elution times over the period of analysis. Duplicates run at six concentrations yield a precision of ±0.09 mg C/L or better. Samples also pass through a UV-Detector (Smartline UV Detector 200, Germany) for determination of the specific ultra-violet absorbance at 254 nm (SUVA), calculated by normalizing the absorbance at this wavelength to the DOM concentration.

### 2.3 Statistical Analyses

Multiple sampling events from the same site have been averaged into one value per site (Suppl. Fig S1; (Aukes et al. 2020)). Principal component analyses (PCA) was performed using *prcomp* in R (R Core Team 2020). Differences in DOM composition across water-body types was assessed via a non-parametric multivariate analysis of variance PERMANOVA using the *adonis2* function in the *vegan* package (Oksanen et al. 2019). Further pairwise comparisons were calculated using a permutational MANOVA via the *EcolUtils* package (Salazar 2020).

## 3 RESULTS

### 3.1 DOM Quantity

Highest DOM concentrations were found in groundwater samples in organic-rich peats from Yellowknife (mean: 97 mg C/L, 1σ: ±40 mg C/L) and ELA (48±22 mg C /L; Figure 2a, Table 1). Lowest concentrations were found in organic-poor groundwater sites (BBW: 0.6±0.4 mg C/L; NRW: 2.1±0.5 mg C /L). The heavily agriculturally-impacted river (GR: 6.1±1.0 mg C /L) contained similar DOM concentrations to pristine Canadian Shield subarctic rivers (YW: 6.6±3.1 mg C /L). Groundwater DOM concentration exhibited a greater difference between ecozones than pond, lake, stream, or river DOM, and were lower in southern areas with mineral strata compared to northern areas with organic-rich deposits. In general, a larger range in DOM concentration was found across groundwater than other water-body types.

**Figure 2:**
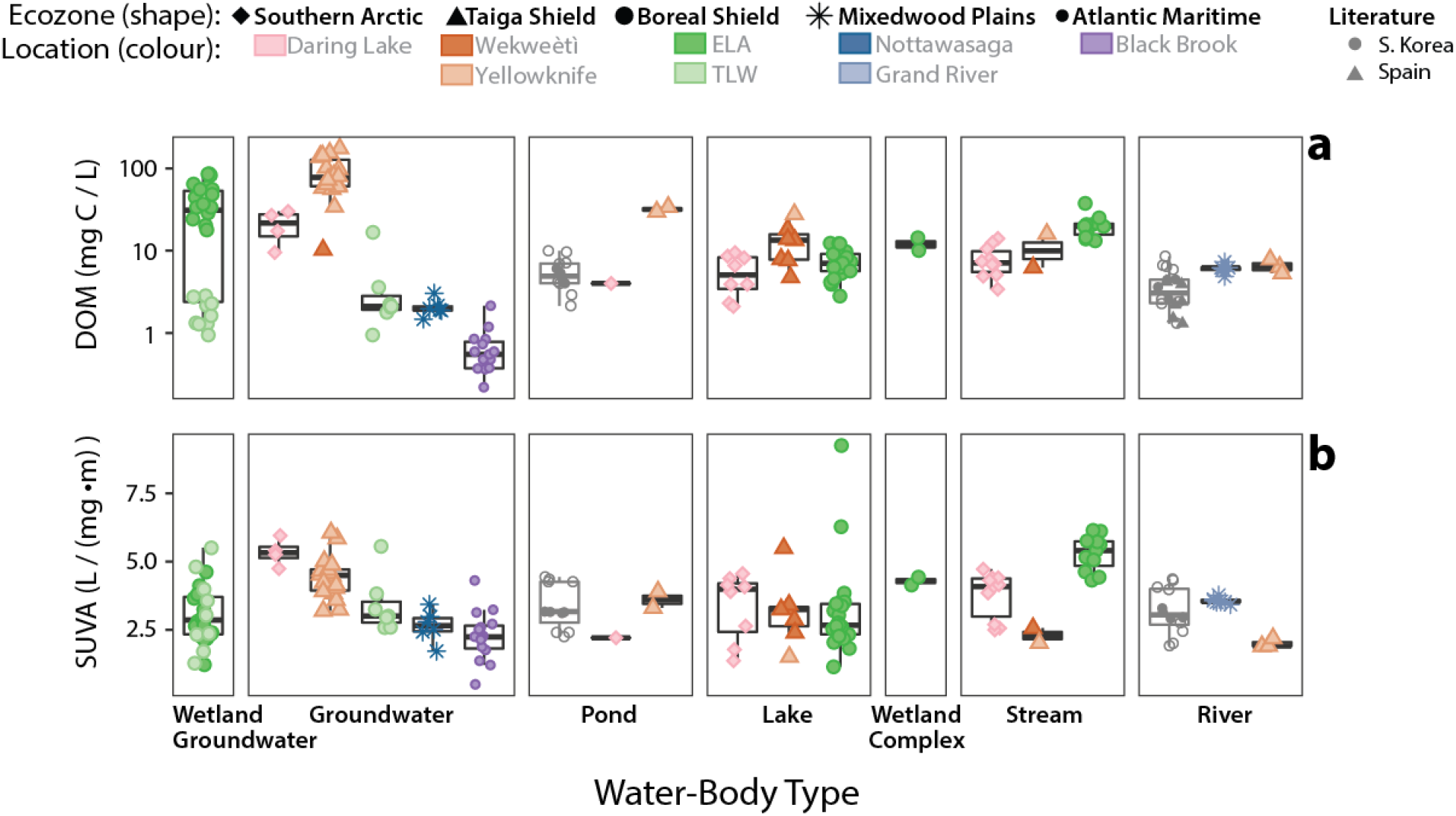
Comparison of dissolved organic matter concentration (DOM; logarithmic y-scale, a) and specific ultra-violet absorbance at 254 nm (SUVA; b) across different water-body types. For each water-body type, different ecozones are grouped together and denoted by different shapes, while separate locations within each ecozone are differentiated by colour. Boxplots for each ecozone in a water-body type indicate the mean value and extend to the first and third quartile, while whiskers represent 1.5x inter-quartile range. Included are literature values that use LC-OCD in South Korea (He et al. 2016) and Spain (Catalán et al. 2017), and are denoted by symbols in grey. Literature values from South Korea include a maximum and minimum value found within the study (open circle). Random scatter is introduced in the x-axis for ease of seeing all data points.

### 3.2 DOM UV-Absorbance

The UV-absorbing capability of DOM can be compared across samples using SUVA values. Lowest SUVA values were measured in agriculturally-impacted groundwater DOM (BBW, NRW) and highest in northern groundwater (DL) and boreal stream DOM (ELA; Figure 2b; Table 1). Unlike samples from ELA, northern ecozone groundwaters (DL, WK, and YW) had higher SUVA values than other water-body types. Highest SUVA values were encountered in three Boreal Shield lakes at depth (~5-10 m below surface) and may represent samples with iron interference (Weishaar et al. 2003) and are therefore not included in subsequent discussion. Overall, differences in groundwater SUVA values exist across ecozones but SUVA values are more similar among other water-body types.

### 3.3 LC-OCD Characterization

Different water-body types within an ecozone contained different proportions of BP. Biopolymers ranged from 1-45% of DOM_hyphl_ but generally contributed less than 15% to DOM_hyphl_ (Figure 3a; Table 1). Streams and lakes contained higher BP proportions than groundwaters. Agricultural groundwater sites contained the lowest BP proportions (Figure 3a). High BP proportions were found in Boreal and Arctic lakes, while moderate BP proportions were found in rivers and were similar to other studies using LC-OCD (He et al. 2016; Catalán et al. 2017) (Figure 3a). Similar BP fraction averages are found across a water-body type compared to different water-body types within an ecozone, and high BP proportions are indicative of lake or stream DOM.

**Figure 3:**
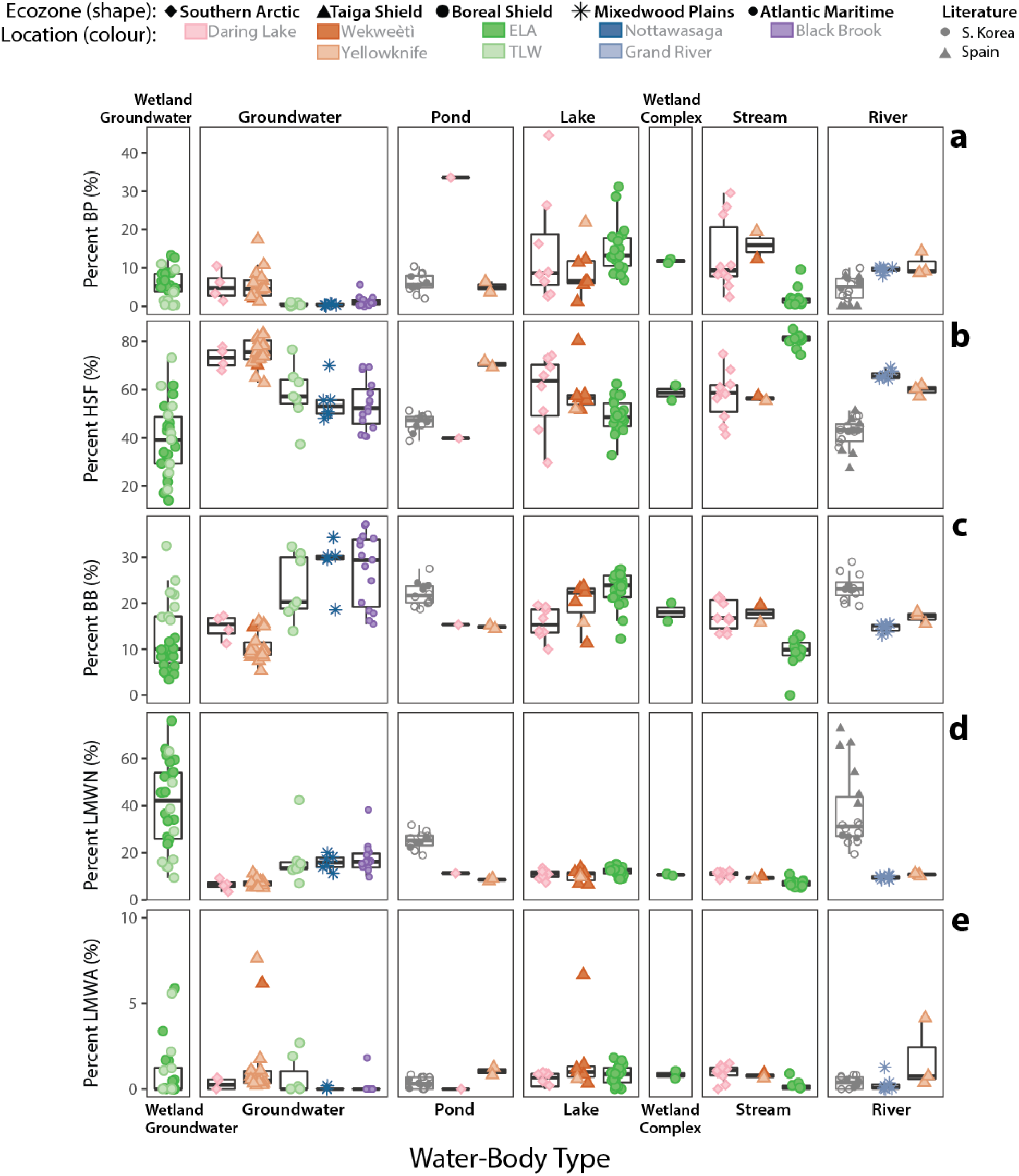
Differences in the composition of DOM based on proportions of LC-OCD groups across water-body types: biopolymers (BP; a), humic substances fraction (HSF; b), building blocks (BB; c), low molecular weight neutrals (LMWN; d), and low molecular weight acids (LMWA; e). For each water-body type, different ecozones are grouped together and denoted by different shapes, while separate locations within each ecozone are differentiated by colour. Boxplots for each ecozone in a water-body type represent the mean value and extend to the first and third quartile, while whiskers represent 1.5x inter-quartile range. Included are literature values that use LC-OCD in South Korea (He et al. 2016) and Spain (Catalán et al. 2017), and are denoted by symbols in grey. South Korea include a maximum and minimum value found within the study (open circle). Random scatter is introduced in the x-axis for ease of seeing all data points.

The HSF comprised the largest proportion, representing up to 85% of DOM_hyphl_ in some environments (Figure 3b). Overall, the HSF proportion ranged from 15-85% with the highest proportions found in Taiga Shield groundwater and Boreal streams (Figure 3). Taiga Shield and Southern Arctic lakes and ponds contained higher HSF proportions than Boreal lakes (Table 1). Taiga Shield rivers had slightly lower proportions than the Mixedwood Plains agriculturally-impacted river. Boreal and agriculturally-impacted (NRW and BBW) groundwaters had the lowest HSF proportions, as well as Boreal wetland groundwater (Table 1; Figure 3).

Smaller molecular weight humics, defined as building blocks (BB), ranged from 5-37% of DOM_hyphl_. High BB proportions were found from southern agricultural groundwater samples, while lowest proportions were observed among a Boreal shield stream and groundwater samples in organic rich peats in YW and ELA (<10%; Figure 3c). Across ecozones, ponds, lakes, streams, and rivers had comparable proportions of BB between 10-30% of DOM_hyphl_.

Proportions of LMWN ranged from 3-76% and were much lower than HSF except among groundwater samples (Figure 3d). DOM from Boreal wetland groundwater sites contained higher LMWN proportions than any other environment (Table 1). Groundwater environments with the lowest HSF proportion had higher LMWN proportions, specifically the agriculturally-impacted sites. Lowest proportions were found from Boreal streams and Arctic groundwater and pond samples. LMWA generally comprised a minor component of total DOM (Figure 3e).

The variability across different sites in LC-OCD defined DOM composition was assessed using PCA. Initially, two principal axes accounted for 60% of the variability within the dataset (Suppl. Fig S2). However, high LMWN proportions from Boreal wetland groundwaters did not allow for a good resolution of other components. For this reason, only these wetland groundwater samples were omitted (n=28) and the PCA was recalculated based on the remaining samples (n=122; Figure 4; Suppl. Table 2). Two principal axes now account for 71% of the variability and illustrates a similarity in DOM composition when comparing water-body types across ecozones. The first principal component (PC) axis represents degraded humics or smaller DOM components, whereas the second PC represents differences in the BP fraction (Figure 4). Distinct compositions are observed comparing lake and groundwater DOM, while rivers and streams contain intermediary compositions between these. Broad variation in groundwater DOM composition resulted from HSF and LMWN or BB proportions, while higher BP proportions were common for ponds, lakes, streams and rivers. Lake DOM composition was also highly variable and mainly characterized by LMWN, BB, and BP. Groundwater DOM from Arctic ecozones (organic soils) grouped separately from other groundwater samples (mineral substrate), but were similar in composition to Boreal streams. Lake DOM can be identified by higher BB and LMWN, lower HSF, and occasionally high BP.

**Figure 4:**
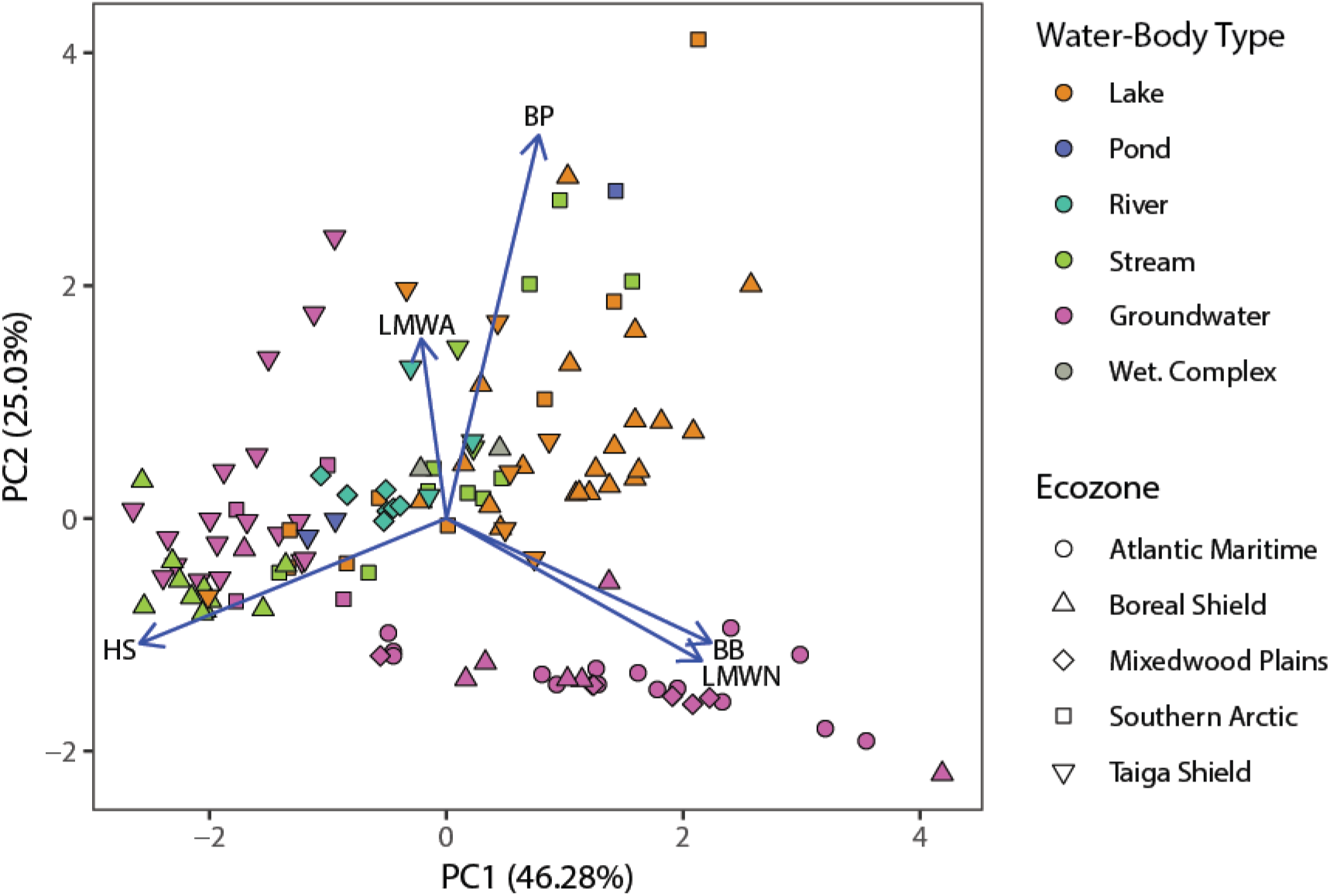
Principal component analyses (PCA) of LC-OCD data (minus wetland groundwater samples) with first and second principal component axes plotted. PCA vectors are included for the proportion of DOM for each LC-OCD group: biopolymers (BP), humic substances fraction (HSF), building blocks (BB), low molecular weight neutrals (LMWN), and low molecular weight acids (LMWA).

Results from the PERMANOVA analyses rejects the null hypothesis and suggests that size-based DOM composition is different between water body types (Table 2). Although the study includes a disproportionate number of water body types per ecozone, results suggest the grouping of DOM compositions by water body type regardless of the vase geographic range encountered in the dataset. For instance, DOM in lakes from boreal shield, southern arctic, and taiga shield all plot together and are significantly different than the composition of river or creek DOM (Table 2).

## 4 DISCUSSION

### 4.1 Can LC-OCD Analyses Differentiate DOM Composition?

This study provided a comprehensive dataset to quantify DOM heterogeneity across a range of environmental conditions that spans the Southern Arctic, with long-cold winters and short-cool summers, to the Mixedwood Plains with warm-wet climates and productive soils. DOM samples are a product of various sources and processes at a single point in time, making it difficult to isolate the relative importance of a specific process or source within a specific ecozone. Differences in both SUVA and size-based properties of DOM fractions are apparent (Figure 2, 3) and result from differences in organic matter sources, catchment characteristics, residence times, and processing history (Curtis and Schindler 1997; Jaffé et al. 2008; Mueller et al. 2012). We find that size-based groupings can identify differences in DOM composition not observed measuring SUVA alone. For instance, ELA and TLW wetland groundwater samples contained similar SUVA values (Figure 2b) but different LC-OCD compositions: a greater proportion of degraded humics at TLW than ELA (Figure 3c; Table 1). The use of various characterization techniques would provide different information on DOM and may be the best way to holistically quantify DOM. Differences in size-based groupings of DOM provide additional information suited to identifying differences in DOM composition that is not provided by UV-based techniques alone.

Although our study design limits the ability to conclusively determine whether DOM composition differed significantly with ecozone, PCA and PERMANOVA analyses suggests a similarity in DOM composition within a water-body type regardless of ecozone (Table 2). Thus, although climate and vegetation differ, DOM composition within a water-body type is comparable across ecozones to be significantly different when compared to other water-body types, offering an interesting avenue for future research.

### 4.2 Similarity in Groundwater DOM

Groupings of similar water-body types in the PCA indicates a commonality of DOM composition across spatial scales. The general conception, especially in groundwater environments, is that processing of DOM results in a loss of heterogeneity. For instance, physiochemical and biological processes were observed to conform DOM composition within a South Carolina soil (Shen et al. 2015). The grouping of groundwater DOM in the PCA indicates various ecozones contain similar mixtures of LC-OCD defined components. Groundwater DOM is differentiated from DOM in other water-body types by the proportion of BP, but can be even further separated by HSF and LMWN to compare organic-rich contributions (ELA and TLW) to agriculturally-impacted sites within mineral strata (BBW and NRW; Figure 3a; Figure 4). Biodegradation and accumulation of DOM in porewaters (Chin et al. 1998) may be responsible for wetland groundwater DOM from Boreal shield sites being easily identified by high proportions of LMWN (Suppl Fig 2). Hence, size-based groupings of groundwater DOM indicates that, while composition can differ among groundwater environments, groundwater contains much different DOM than other water-body types.

### 4.3 Trends in DOM Composition across Water-Body Types

Photolysis and *in-situ* production are two processes that occur among surface waters that alter the composition of DOM. Exposure of DOM to sunlight and photochemical transformations of humics results in smaller molecules and increased proportions of BB (Tercero Espinoza et al. 2009; He et al. 2016). This is observed in the dataset by a shift from PCA groupings with high BP (indicative of lakes, ponds, and streams) towards higher BB and LMWN suggestive of exposure to sunlight (Figure 4). High BP in lakes, rivers, and most streams indicate high molecular weight polysaccharides or proteins that form during *in-situ* production or microbial processing of DOM (Ciputra et al. 2010; Huber et al. 2011; Catalán et al. 2017).

Rivers and streams can act as conduits of DOM transport and internal producers and processors of DOM (Creed et al. 2015), forming a DOM composition intermediary between lake and groundwater end-members. Further, stream DOM composition can identify DOM source as some samples plot closely with either groundwater or lake DOM (Figure 4). Thus, size-based DOM groupings can be used to evaluate the importance of autochthonous versus groundwater sources of DOM in rivers and streams.

Along a hypothetical flow path, terrestrial groundwater DOM is mobilized into surrounding ponds and lakes, and eventually exported from the watershed via rivers or streams. Changes to LC-OCD components, concurrent with decreases to overall DOM concentration, are found along this hypothetical flow with the loss of high-HSF in groundwaters as LMW and BP proportions increase (Figure 3, 4). These changes are in agreement with watershed-scale observations of DOM composition change along an aquatic continuum, shifting from relatively high-molecular weight groundwater components to smaller, more degraded, and more aliphatic components (Kellerman et al. 2014; Hutchins et al. 2017). The similarity in DOM composition within water-body types across different ecoregions (Figure 4) indicates the processes responsible for altering DOM composition may be similar regardless of surrounding vegetation or climate.

### 4.4 Implications of Different Size-Based Groupings of DOM

Differences in DOM composition are associated with water-body type, hence shifts or changes to the hydrological regime, such as recent increases to terrestrial-derived DOM, can alter DOM composition and its effect upon the surrounding environment (Creed et al. 2018). Size-exclusion DOM analysis provides a quantitative measure of DOM composition variability across aquatic environments, and may help identify lakes experiencing stronger terrestrial influences, such as higher HSF and lower BP. In particular, a shift towards heavier fall rains in the Northwest Territories, Canada (Spence et al. 2011), can enhance terrestrial DOM contributions to surrounding surface waters, which may result in increased pre-treatment costs (Ritson et al. 2014) and higher DBP concentrations (Awad et al. 2016). Implementation of this size-based grouping scheme provides a quantitative tool to measure DOM composition while acknowledging the inherent heterogeneity when attempting to characterize the complex mixtures characteristic of naturally occurring DOM. The ability to characterize and compare specific DOM compositions is important to better predict, plan, and adapt water treatment methods for these climate-induced changes.

## ACKNOWLEDGEMENTS

Thanks to the Environmental Geochemistry Laboratory at the University of Waterloo for assistance with sample collection and processing. We thank Michael English, Roy Judas, and Jordan Reid for field assistance in collecting samples in the Northwest Territories, and the Community of Wekweètì and the Wek’èezhìi Land & Water Board for their support during our sampling events. Staff of the Fredericton Research and Development Centre, Agriculture and Agri-Food Canada, collected samples from the Black Brook Watershed, New Brunswick. This work benefited from the assistance of Monica Tudorancea with operation of the LC-OCD instrument. We would also like to thank four anonymous reviewers for their suggestions and comments. Funding was provided by a Natural Science and Engineering Research Council (NSERC) of Canada grant to S. L. Schiff, and an Ontario Graduate Scholarship, University of Waterloo Scholarship, and International Association of GeoChemistry Research Award (IAGC) to P. J. K. Aukes. Additional financial support was provided to J. Spoelstra by Environment and Climate Change Canada and the Ontario Ministry of Agriculture, Food, and Rural Affairs (OMAFRA). The authors declare no conflicts of interest related to the study. All data can be found in the Supplementary Information, while code used to generate figure and statistical analysis used in this manuscript is available at github.com/paukes/canada-ecozone-DOM.

## SUPPLEMENTARY INFORMATION

### LC-OCD Methodology

Dissolved organic matter characterization using liquid chromatography – organic carbon detection (LC-OCD) was performed in the Department of Civil and Environmental Engineering, University of Waterloo. A detailed description of the instrument, setup, and analysis can be found in Huber et al. (2011). First, samples are diluted to a DOM concentration between 1 – 5 mg C/L to obtain an optimal chromatogram to analyse. Dilution factors were used to correct concentrations for comparison. The sample is added to a mobile phase (phosphate buffer exposed to UV-irradiation) and passed through an in-line 0.45μm filter and into either a by-pass or the size-exclusion column (Toyopearl HW-50S (Tosoh, Japan)). The by-pass allows for overall determination of the dissolved organic carbon concentration, as well as the overall UV-absorbance at 254 nm. The novel organic carbon detector (OCD) is comprised of a thin film UV-reactor (Gräntzel thin-film reactor, DOC-Labor, Karlsruhe, Germany). As the acidified sample enters the OCD, it is thinly spread over a UV-lamp, allowing irradiation to oxidize organic carbon into CO_2_. The CO_2_ is then measured with a highly-sensitive infrared detector, where the integration of the peak area provides the total DOC concentration. Sample that is not passed through the by-pass enters the size-exclusion column (SEC) where it is divided into different fractions based on nominal molecular weight and is continuously analyzed by the UV detector and OCD, producing OCD and UV chromatograms.

### LC-OCD Molecular Groupings

Chromatograms are analyzed using customized software (ChromCALC, DOC-LABOR, Karlsruhe, Germany) which calculates the concentration of each ‘size grouping’ by integrating areas based on specific elution times. First, the overall DOC concentration is determined from the bypass peak. Second, the hydrophilic portion (sample that elutes from the SEC) is subdivided into five categories based upon elution time: biopolymers (BP), humic substances fraction (HSF; which includes both humic and fulvic acids), building blocks (BB), and low-molecular-weight neutrals (LMW-N) and acids (LMW-A).

Grouping names and boundaries of each fraction are arbitrarily defined based on elution times and characteristics as determined by Huber et al. (2011). Briefly, the groupings are (from largest to smallest):

- Biopolymers: >10 kDa; polysaccharides, proteins, or amino sugars
- Humic substances fraction: 100 – 1200 Da; a heterogeneous mixture of large, complex molecules that elute at similar times as the IHSS humic standards
- Building blocks: degraded humic substances with humic-like characteristics, but of lower molecular weight
- LMW acid and LMW neutrals: contain monoprotic acids, amino sugars, ketones, and aldehydes Finally, the software determines a ‘hydrophobic’ parameter quantified as the residual between the sum of all measured fractions and the overall concentration of DOC (determined from the by-pass). As the difference between the overall DOM concentration and the sum of the five eluted fractions. As the hydrophilic component comprised 90±9% across all DOM samples, group percentages were normalized to the sum of the eluted components.

## SUPPLEMENTARY TABLES

**Table S1:**
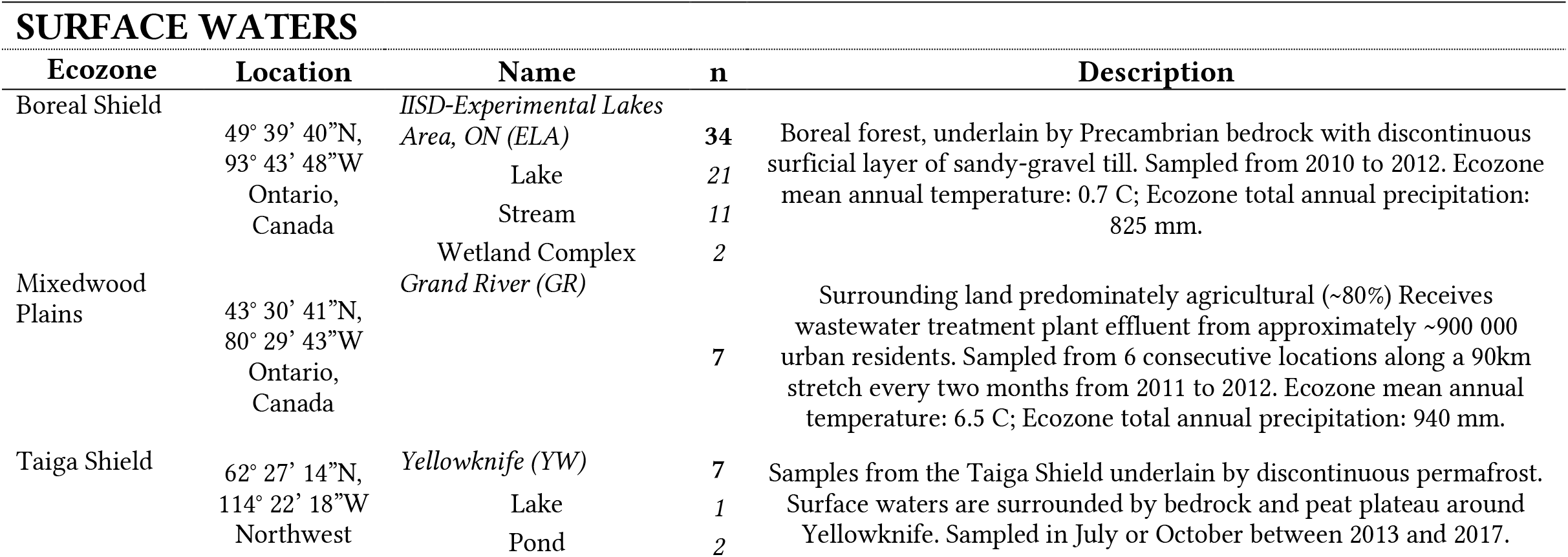
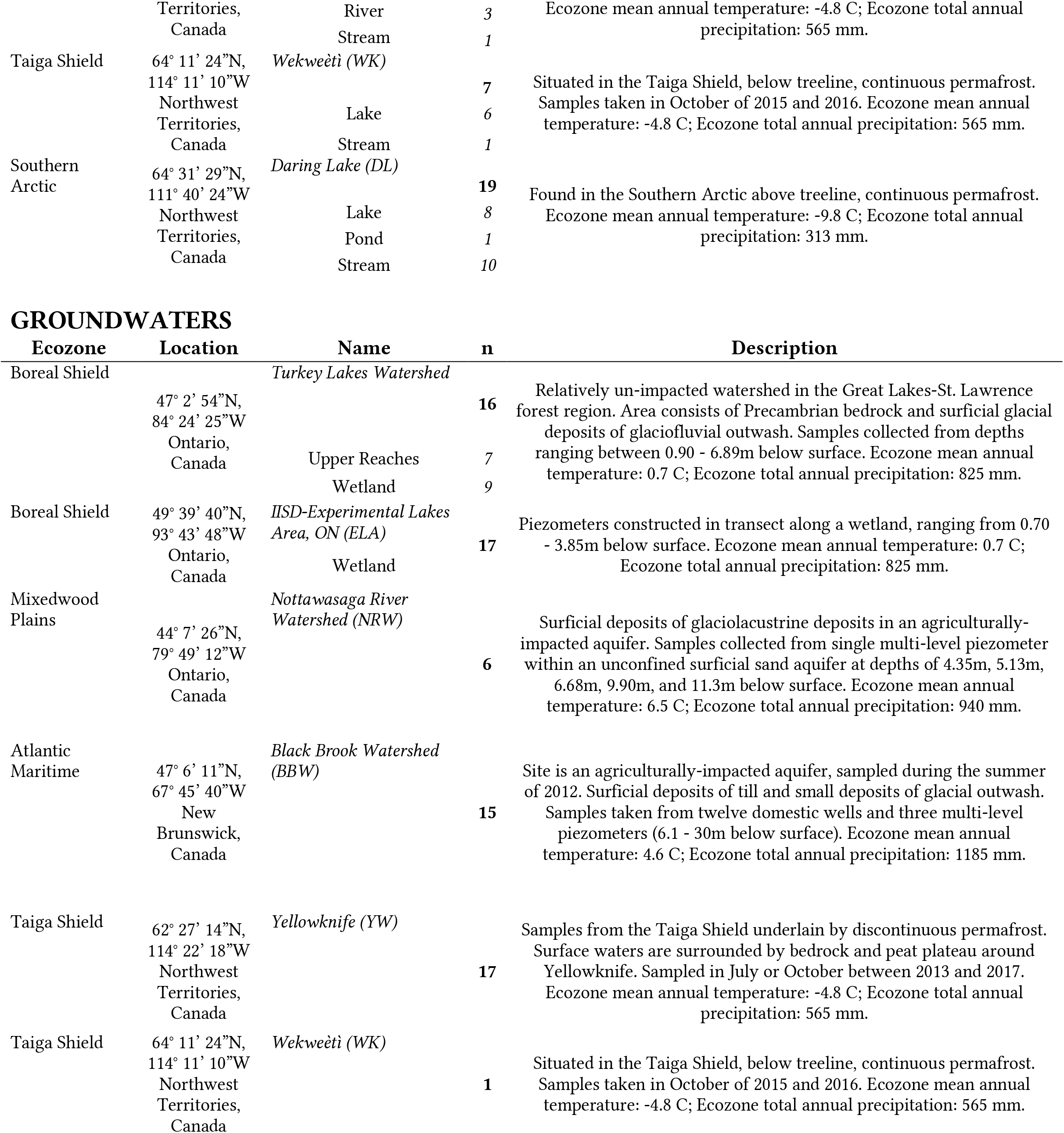
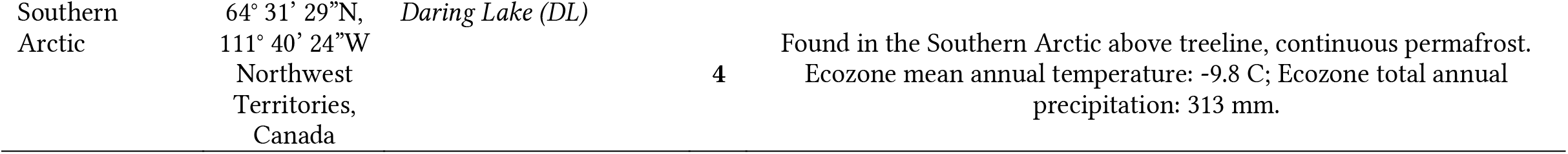
Description of sampling sites. The number of samples represents the number of distinct sites sampled in that water-body for that ecozone. Mean annual air temperatures and total annual precipitation taken from Marshall et al. (1999).

**Table S2:**
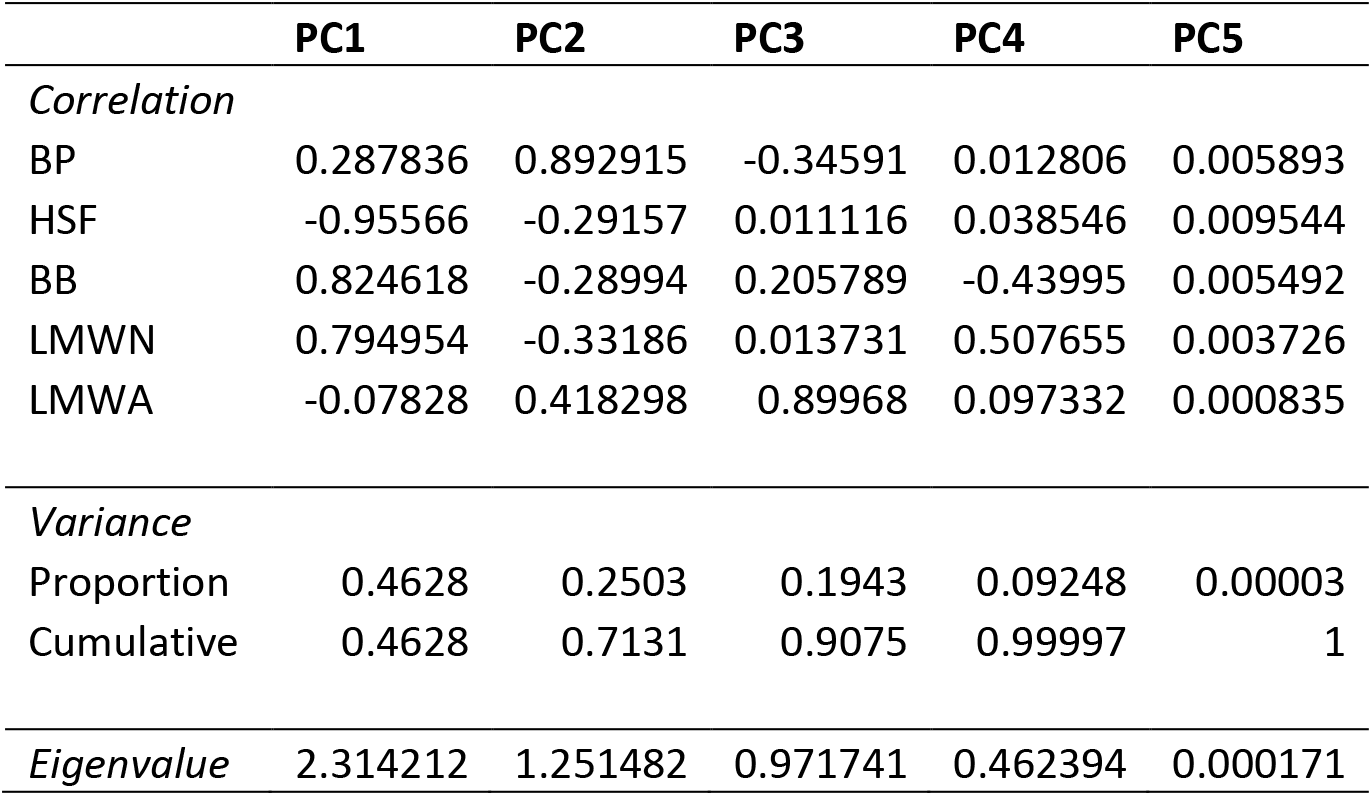
Summary statistics for the principal component analysis used in Figure 4. Included are the LC-OCD variables and their correlation with each principal component (PC) axis, variance explained by each PC, and eigenvalues for each PC. All code used to generate these results can be found on github.com/paukes/canada-ecozone-DOM.

## SUPPLEMENTARY FIGURES

**Figure S1:**
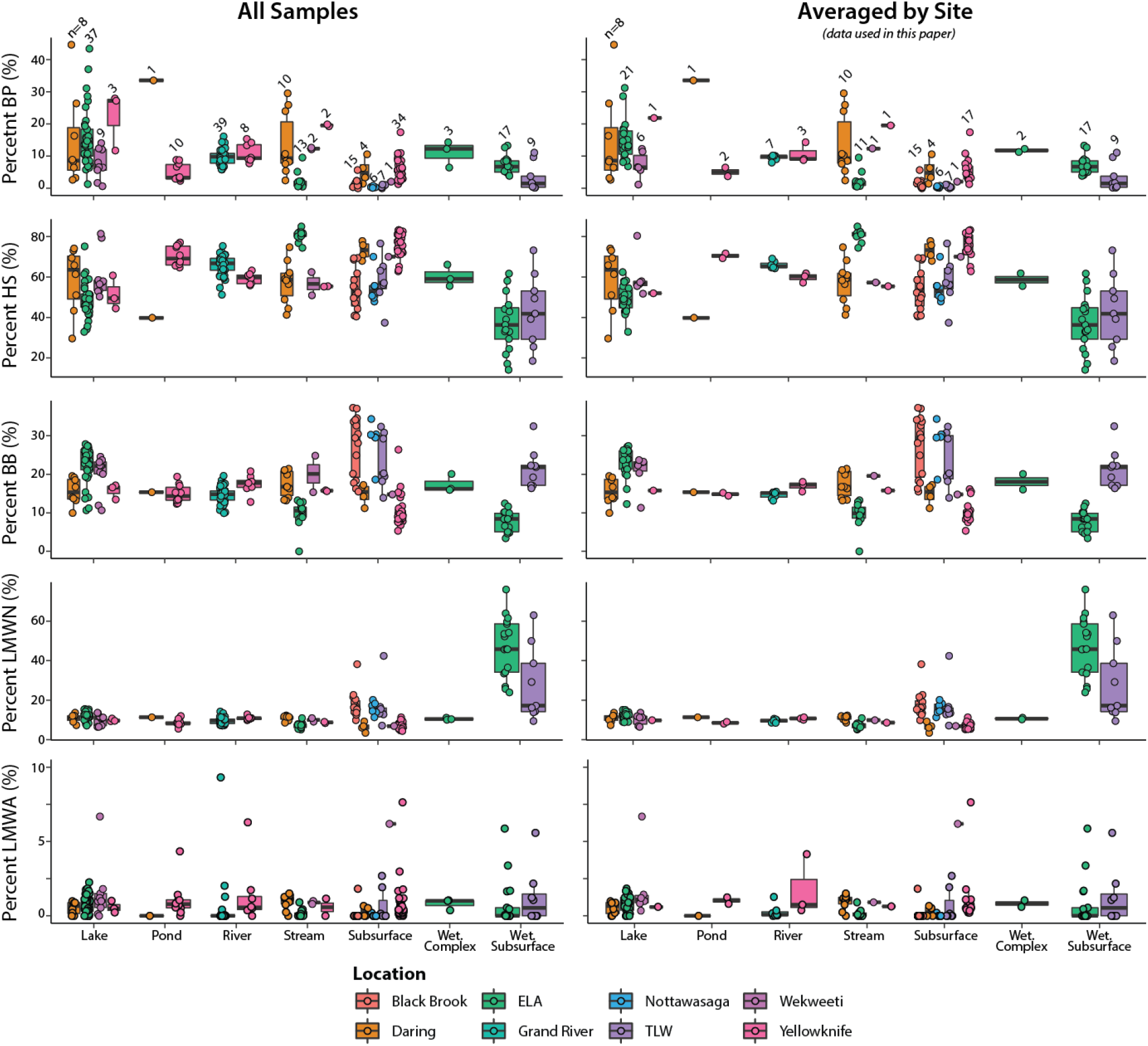
Comparison of LC-OCD fractions for each water-body type between all sampling events (left graph; including numerous sampling events at the same site) and averaged-sampling events per site (right panel; one averaged number per unique site) for different locations (denoted by colour). A boxplot of the data is overlain by the actual data points for each site (with random scatter on the x-axis to improve visibility of all points) and the total number of points is included as a black number above each boxplot. Each boxplot represents the mean with 25th and 75th percentiles, with whiskers that extend up to 1.5x the inter-quartile range.

**Figure S2:**
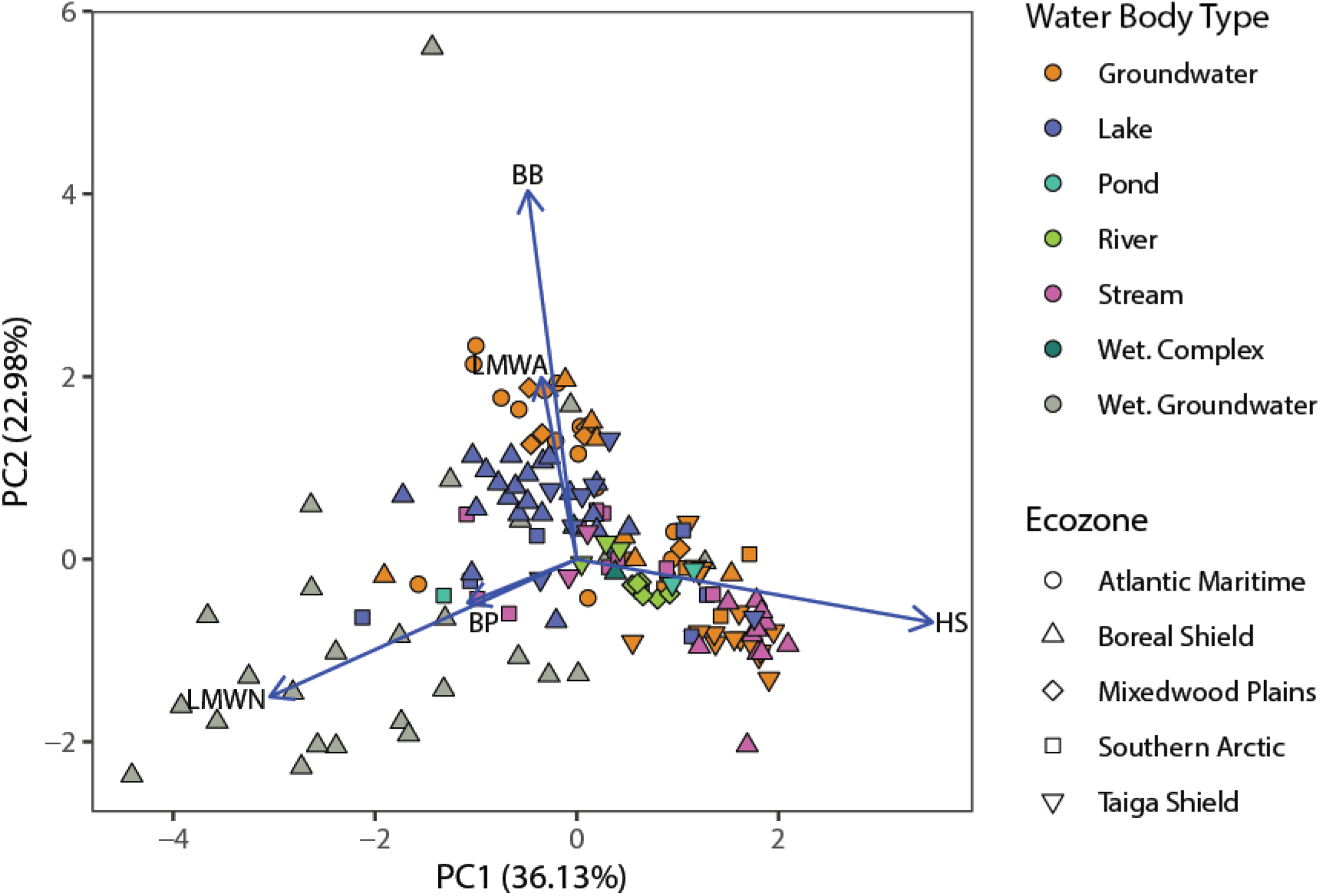
Original principal component analysis (PCA) that included wetland groundwater samples (pink cross symbol) with high DOM and high LMWN proportions.

