## Supplemental Information for "Size-Based Characterization of Freshwater Dissolved Organic Matter finds Similarities within a Water Body Type across Different Canadian Ecozones"

#### SUPPLEMENTARY TABLES

*Table S1: Description of sampling sites. The number of samples represents the number of distinct sites sampled in that water-body for that ecozone. Mean annual air temperatures and total annual precipitation taken from Marshall et al. (1999).*

**SURFACE WATERS**

| Ecozone | Location | Name | n | Description |
|---|---|---|---|---|
| Boreal Shield | 49° 39' 40"N, 93° 43' 48"W Ontario, Canada | *IISD-Experimental Lakes Area, ON (ELA)* | **34** | Boreal forest, underlain by Precambrian bedrock with discontinuous surficial layer of sandy-gravel till. Sampled from 2010 to 2012. Ecozone mean annual temperature: 0.7 C; Ecozone total annual precipitation: 825 mm. |
| | | Lake | *21* | |
| | | Stream | *11* | |
| | | Wetland Complex | *2* | |
| Mixedwood Plains | 43° 30' 41"N, 80° 29' 43"W Ontario, Canada | *Grand River (GR)* | **7** | Surrounding land predominately agricultural (~80%) Receives wastewater treatment plant effluent from approximately ~900 000 urban residents. Sampled from 6 consecutive locations along a 90km stretch every two months from 2011 to 2012. Ecozone mean annual temperature: 6.5 C; Ecozone total annual precipitation: 940 mm. |
| Taiga Shield | 62° 27' 14"N, 114° 22' 18"W Northwest | *Yellowknife (YW)* | **7** | Samples from the Taiga Shield underlain by discontinuous permafrost. Surface waters are surrounded by bedrock and peat plateau around Yellowknife. Sampled in July or October between 2013 and 2017. |
| | | Lake | *1* | |
| | | Pond | *2* | |

| | | | | |
|---|---|---|---|---|
| | Territories, Canada | River | *3* | Ecozone mean annual temperature: -4.8 C; Ecozone total annual precipitation: 565 mm. |
| | | Stream | *1* | |
| Taiga Shield | 64° 11’ 24”N, 114° 11’ 10”W Northwest Territories, Canada | *Wekweètì (WK)* | **7** | Situated in the Taiga Shield, below treeline, continuous permafrost. Samples taken in October of 2015 and 2016. Ecozone mean annual temperature: -4.8 C; Ecozone total annual precipitation: 565 mm. |
| | | Lake | *6* | |
| | | Stream | *1* | |
| Southern Arctic | 64° 31’ 29”N, 111° 40’ 24”W Northwest Territories, Canada | *Daring Lake (DL)* | **19** | Found in the Southern Arctic above treeline, continuous permafrost. Ecozone mean annual temperature: -9.8 C; Ecozone total annual precipitation: 313 mm. |
| | | Lake | *8* | |
| | | Pond | *1* | |
| | | Stream | *10* | |

### GROUNDWATERS

| Ecozone | Location | Name | n | Description |
|---|---|---|---|---|
| Boreal Shield | 47° 2’ 54”N, 84° 24’ 25”W Ontario, Canada | *Turkey Lakes Watershed* | **16** | Relatively un-impacted watershed in the Great Lakes-St. Lawrence forest region. Area consists of Precambrian bedrock and surficial glacial deposits of glaciofluvial outwash. Samples collected from depths ranging between 0.90 - 6.89m below surface. Ecozone mean annual temperature: 0.7 C; Ecozone total annual precipitation: 825 mm. |
| | | Upper Reaches | *7* | |
| | | Wetland | *9* | |
| Boreal Shield | 49° 39’ 40”N, 93° 43’ 48”W Ontario, Canada | *IISD-Experimental Lakes Area, ON (ELA)* | **17** | Piezometers constructed in transect along a wetland, ranging from 0.70 - 3.85m below surface. Ecozone mean annual temperature: 0.7 C; Ecozone total annual precipitation: 825 mm. |
| | | Wetland | | |
| Mixedwood Plains | 44° 7’ 26”N, 79° 49’ 12”W Ontario, Canada | *Nottawasaga River Watershed (NRW)* | **6** | Surficial deposits of glaciolacustrine deposits in an agriculturally-impacted aquifer. Samples collected from single multi-level piezometer within an unconfined surficial sand aquifer at depths of 4.35m, 5.13m, 6.68m, 9.90m, and 11.3m below surface. Ecozone mean annual temperature: 6.5 C; Ecozone total annual precipitation: 940 mm. |
| Atlantic Maritime | 47° 6’ 11”N, 67° 45’ 40”W New Brunswick, Canada | *Black Brook Watershed (BBW)* | **15** | Site is an agriculturally-impacted aquifer, sampled during the summer of 2012. Surficial deposits of till and small deposits of glacial outwash. Samples taken from twelve domestic wells and three multi-level piezometers (6.1 - 30m below surface). Ecozone mean annual temperature: 4.6 C; Ecozone total annual precipitation: 1185 mm. |
| Taiga Shield | 62° 27’ 14”N, 114° 22’ 18”W Northwest Territories, Canada | *Yellowknife (YW)* | **17** | Samples from the Taiga Shield underlain by discontinuous permafrost. Surface waters are surrounded by bedrock and peat plateau around Yellowknife. Sampled in July or October between 2013 and 2017. Ecozone mean annual temperature: -4.8 C; Ecozone total annual precipitation: 565 mm. |
| Taiga Shield | 64° 11’ 24”N, 114° 11’ 10”W Northwest Territories, Canada | *Wekweètì (WK)* | **1** | Situated in the Taiga Shield, below treeline, continuous permafrost. Samples taken in October of 2015 and 2016. Ecozone mean annual temperature: -4.8 C; Ecozone total annual precipitation: 565 mm. |

| | | | | |
|---|---|---|---|---|
| Southern Arctic | 64° 31' 29"N, 111° 40' 24"W Northwest Territories, Canada | *Daring Lake (DL)* | **4** | Found in the Southern Arctic above treeline, continuous permafrost. Ecozone mean annual temperature: -9.8 C; Ecozone total annual precipitation: 313 mm. |

*Table S2: Summary statistics for the principal component analysis used in Figure 4. Included are the LC-OCD variables and their correlation with each principal component (PC) axis, variance explained by each PC, and eigenvalues for each PC. All code used to generate these results can be found on github.com/paukes/canada-ecozone-DOM.*

| | **PC1** | **PC2** | **PC3** | **PC4** | **PC5** |
|---|---|---|---|---|---|
| *Correlation* | | | | | |
| BP | 0.287836 | 0.892915 | -0.34591 | 0.012806 | 0.005893 |
| HSF | -0.95566 | -0.29157 | 0.011116 | 0.038546 | 0.009544 |
| BB | 0.824618 | -0.28994 | 0.205789 | -0.43995 | 0.005492 |
| LMWN | 0.794954 | -0.33186 | 0.013731 | 0.507655 | 0.003726 |
| LMWA | -0.07828 | 0.418298 | 0.89968 | 0.097332 | 0.000835 |
| *Variance* | | | | | |
| Proportion | 0.4628 | 0.2503 | 0.1943 | 0.09248 | 0.00003 |
| Cumulative | 0.4628 | 0.7131 | 0.9075 | 0.99997 | 1 |
| *Eigenvalue* | 2.314212 | 1.251482 | 0.971741 | 0.462394 | 0.000171 |

#### SUPPLEMENTARY FIGURES

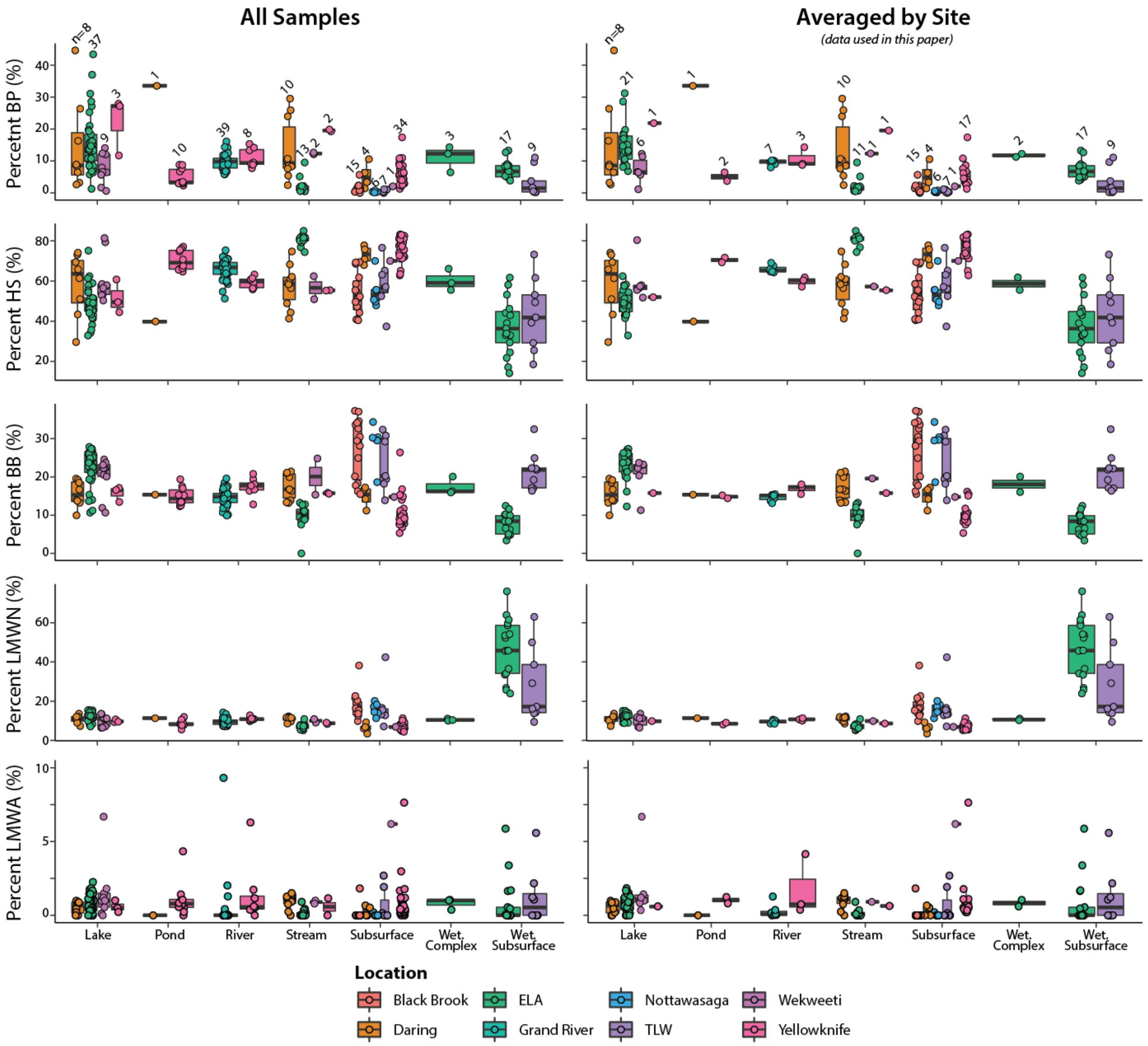


*Figure S1*: Comparison of LC-OCD fractions for each water-body type between all sampling events (left graph; including numerous sampling events at the same site) and averaged-sampling events per site (right panel; one averaged number per unique site) for different locations (denoted by colour). A boxplot of the data is overlain by the actual data points for each site (with random scatter on the x-axis to improve visibility of all points) and the total number of points is included as a black number above each boxplot. Each boxplot represents the mean with 25th and 75th percentiles, with whiskers that extend up to 1.5x the inter-quartile range.

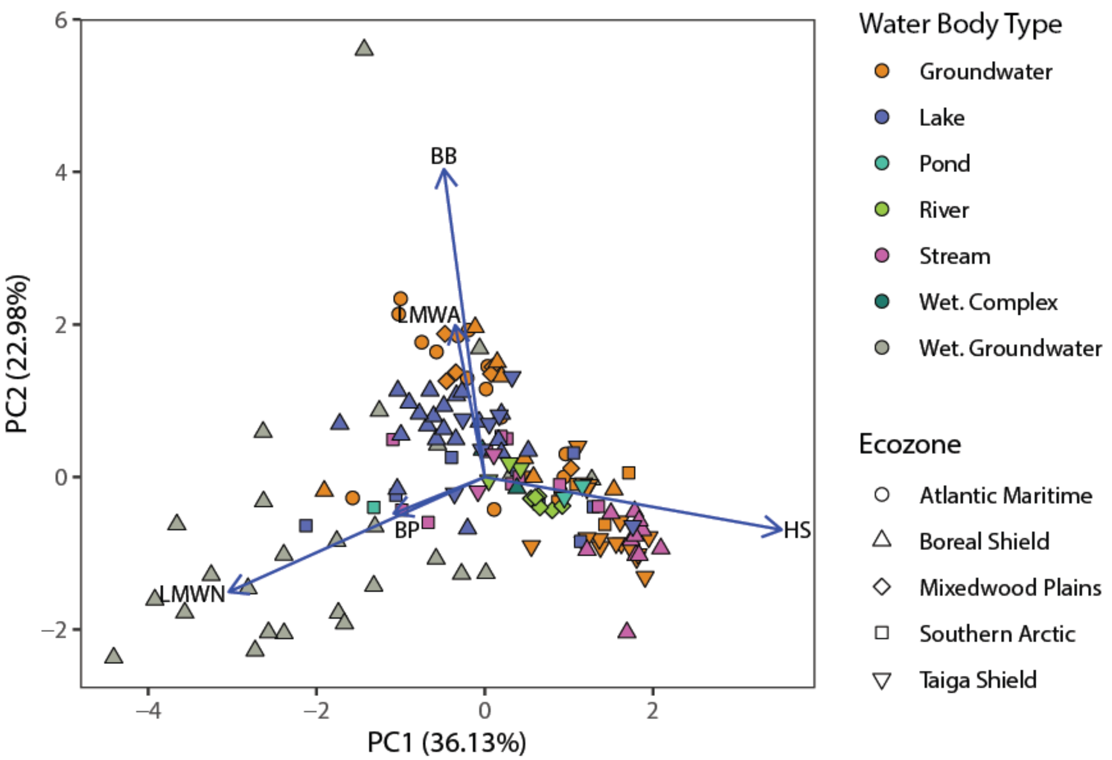


*Figure S2*: Original principal component analysis (PCA) that included wetland groundwater samples (pink cross symbol) with high DOM and high LMWN proportions.
